# Gaseous Signaling Compounds (Hydrogen Sulfide and Nitric Oxide) and Their Relative Roles in Affecting Anaerobic HeLa 229 Cell Viability

**DOI:** 10.1101/2021.05.24.445475

**Authors:** Balbina J. Plotkin, Ira M. Sigar, Amber Kaminski

**Affiliations:** Department of Microbiology and Immunology, College of Graduate Studies, Midwestern University, Downers Grove, IL, USA; (I.S.M.); (A.K.)

**Keywords:** cysteine, hydrogen sulfide, arginine, xanthine, nitric oxide, hypoxia inducible factor (HIF), anoxia

## Abstract

Metabolic pathways supporting long-term anaerobic cell viability have not been identified. The effect NO and H_2_S pathway effectors have on HeLa 229 cell viability was measured after 10 days anaerobic incubation. The addition of arginine or xanthine (NO pathway precursors) consistently increased HeLa cell viability by 13.1- and 4.4-fold, respectively. Allopurinol, a xanthine oxidase inhibitor, also increased viability, as compared to control levels. In contrast, inhibition of iNOS by 1400W increased cell viability by 79-fold. Regarding the H_2_S pathway, precursor cysteine enhanced viability by 9.8-fold with the greatest number of viable cells measured in response to the presence of a H_2_S donor (GYY4137), or an inhibitor of glutathione synthesis, propargylglycine (40- and 85-fold, respectively). These results demonstrate that the constitutive level of cell viability after extended (10 days) growth without oxygen can be modulated by affecting NO or H_2_S generating pathways.

## 1 Introduction

A majority of studies on tumor cell physiology are performed under carbon dioxide enriched atmospheric oxygen conditions. However, this oxygen-rich environment is not physiologically relevant. *In vivo*, the ecosystem ranges from completely lacking in oxygen (anoxic/anaerobic) to hypoxic (less than 13%) (Cojoc et al., 2015; Ivanovic, 2009). While there has been a push towards characterizing cell metabolism under physiologically relevant oxygen levels (physioxia), cell metabolism as it relates to growth under strict anoxic microenvironmental conditions has not been studied. To date, assays performed in an anaerobic macroenvironment are typically initiated using media containing atmospheric oxygen levels that deplete over time; thus, yielding incubation conditions that cover an oxygen concentration spectrum, the nadir of which is not measured (Vlaski-Lafarge et al., 2020).

Characterization of anaerobic cell growth is crucial, particularly since like stem cells in their niches, solid tumors also contain areas lacking an oxygen supply (anaerobic) (Cummins and Tangney, 2013; Gronroos et al., 2014; Visvader and Lindeman, 2008). It is essential to characterize this aspect of cancer cell metabolism to fully understand metastasis and drug resistance. Until recently, the ability to study long-term anoxic cell growth has been elusive, due to an inability to culture cells *ex vivo* for extended periods of time in the absence of oxygen. We pioneered a system that mimics the lack of oxygen in tumor centers from where cancer stem cells, and potentially metastatic cells, are reported to originate. *In vitro*support of the role anaerobic rewiring may enhance tumorigenesis is the finding that HeLa cells grown anaerobically shift their cytokine secretome expression to one that is pro-angiogenic (Plotkin et al., 2018b). The pathways that play a role in long-term anaerobic cell viability have not been identified. The gaseous intercellular transmitters e.g., nitric oxide (NO) and hydrogen sulfide (H_2_S) serve critical roles in tumor microenvironments and acquired chemotherapy resistance (Chen et al., 2019; Cheng et al., 2014; De Vries and Schröder, 2002; Szabo et al., 2013). Since bacteria, which undergo anaerobic respiration, use NO and H_2_S as components of their respiratory pathways, we focused on determining whether NO and H_2_S generating pathways play a role in sustaining anaerobic cancer cell viability, as an index of metabolism (Pal et al., 2018; Plotkin et al., 2015; Stoimenova et al., 2007; Trageser and Unden, 1989).

## 2 Materials and Methods

### Reagents

Effectors for nitric oxide synthesis used in experiments were S-nitroso-N-acetylpenicillamine (SNAP; 50μM; Cayman Chemical), arginine (10mM; Sigma), N omega-nitro-L-arginine methyl ester hydrochloride (L-NAME; 0.15mM; Sigma), N-([3-(aminomethyl)phenyl] methyl) ethanimidamide dihydrochloride (1400W; 100μM; Sigma), xanthine (4×10^-5^M; Sigma), nitrite (50, 100, 500mM; Sigma), and allopurinol (200μM; Sigma). Effectors for hydrogen sulfide synthesis used in experiments were cysteine (0.4mM; Sigma), JK-2 (300μM; Sigma), P-(4-Methoxyphenyl)-P-4-morpholinyl-phosphinodithioic acid (GYY4137; 400μM; Cayman Chemical), pyridoxal phosphate (PLP; 2μM; Sigma), and DL-propargylglycine (PAG; 0.5mM; Sigma).

### Anoxic Cell Culture

Previous studies show that regardless of cell line screened, all can replicate in the absence of oxygen (Plotkin et al., 2018a). For this initial study, to assess the roles various pathways and their mediators play in supporting anaerobic cell viability, HeLa 229 cells were used based on their replication rate and consistency as the model cell line for initial determinations of anaerobic metabolic changes. HeLa 229 cells were grown to 80% confluence (2.24 x 10^5^ cells/well) in 24 well plates (normoxic conditions, 5% CO_2_ in air; 10% FBS, 0.05mg/ml gentamicin (Hyclone); high glucose (4.5 g/L) DMEM medium with glutamine and pyruvate (584 and 110 mg/L, respectively; Cellgro) for 24h. These cells were then transferred to an anaerobic chamber (Whitley A35, anaerobic gas mixture: H_2_, CO_2_, N_2_; 37°C), and medium replaced with degassed low glucose (1 g/L) homologous medium without gentamicin, as previously described (Plotkin et al., 2018a). The lack of oxygen (0%) in the degassed medium and culture plate wells were confirmed by oxygen electrode measurements (Microelectrodes, Inc. oxygen probe; Pod-Vu software).

### Nitric Oxide and Hydrogen Sulfide Pathway Constituents and Selected Inhibitors

Nitric oxide and hydrogen sulfide producing pathways could participate in supporting anaerobic mitochondrial respiration. To test this hypothesis, precursors, inhibitors, and agonists of these pathways were tested for their effect on cell viability and hypoxia inducible factor (HIF) expression. Test chemicals were dissolved in degassed medium under anaerobic conditions. Upon transferring the cells to the anaerobic chamber (Day 0), aerobic DMEM was removed by aspiration and replaced with anaerobic degassed DMEM alone, or containing chemical supplement indicated. At Day 10, the total number of viable cells per condition were determined by trypan blue dye exclusion (Countess™ II Automated Cell Counter) and HIF expression measured (see below). Controls for all tests consisted of cells grown in anaerobic medium alone. Medium with DMSO (0.05%) was used as the control for chemicals requiring DMSO for solution preparation (allopurinol, 1400W, and GYY4137).

### HIF-1α Analysis

HIF expression for each growth condition was determined. Immediately, post-viability measurements were taken, plate well contents (on ice) from each condition were pooled for six wells (4°C), then centrifuged (2000 RPM, 5 min, 4°C). Supernatants were removed. HIF-1α protein was extracted from cell pellets and confirmed using Human Simple Step ELISA kit (Abcam) according to manufacturer’s instructions.

### Data Analysis

Data were analyzed (GraphPad Prism) by two-way ANOVA (*p*≤ 0.05). Where appropriate, Tukey post-hoc tests were performed. Significant points (*p* < 0.05) are designated with an asterisk and standard error of the mean (SEM) are included on all graphs.

## 3 Results and Discussion

In solid tumors, there are intra-tumor microenvironments that are anaerobic. As previously reported, replicating HeLa cells at 10 days anaerobic incubation produce reactive oxygen species (ROS), as detected by DCFDA and CellRox Green fluorescence, lack expression of the autophagy marker LC3B and accumulate MitoTracker Red FM (Plotkin et al., 2018a). Two important pathways that result in NO release are the oxidation of L-arginine by nitric oxide synthase (NOS), and the nitrite reductase pathway (xanthine oxidase, XO) that can utilize either xanthine, or nitrite as a precursor (Cantu-Medellin and Kelley, 2013; Damacena-Angelis et al., 2017). In bacteria and plants, nitrite can fuel ATP production via the mitochondrial electron transport chain with NO as the terminal electron acceptor (Gladwin et al., 2005; Siva Raju et al., 1997; Stoimenova et al., 2007). In mammalian cells, XO oxidizes either xanthine or nitrite to form NO. The ability of nitric oxide generating pathway constituents to support long-term (10 days) anaerobic cell viability was determined (Fig. 1A; 1B). Of the three fundamental precursors, the addition of xanthine and arginine resulted in a reproducible pattern of increased cell viability by 13- and 4.4-fold, respectively, as compared to supplement-free control, with no associated increase in HIF expression (Fig. 1C). Allopurinol, an inhibitor of xanthine oxidase, resulted in only a 3-fold change in cell viability, while inhibition of the constitutive NOS (cNOS) by L-NAME had no effect on cell viability as compared to control. In contrast, the combination of allopurinol and L-NAME enhanced cell viability by 97-fold, as did the addition of the iNOS inhibitor 1400W alone, and when combined with L-NAME (80-fold and 103-fold increase, respectively). All other supplements tested either had no effect on constitutive anaerobic cell viability, or increased viability except for SNAP, a nitric oxide donor, which depressed viability levels to 0.6-fold of control. Thus, with respect to NO pathways, inhibition of NO production via 1400W and allopurinol (xanthine oxidase) had the most robust effect on increasing cell viability. Another interesting finding relative to anaerobic metabolism that differentiates from what is reported for cell metabolism under hypoxic conditions is that the change in viability appears disconnected from HIF expression. Except for the addition of nitrite (alone or in combination), HIF protein levels were depressed as compared to medium control. Of the supplements tested, only the presence of nitrite significantly enhanced HIF expression, as would be predicted based on reports that nitrite helps stabilize HIF (Diwakar and Ravindranath, 2007) (Fig 1C).

**Fig. 1.**
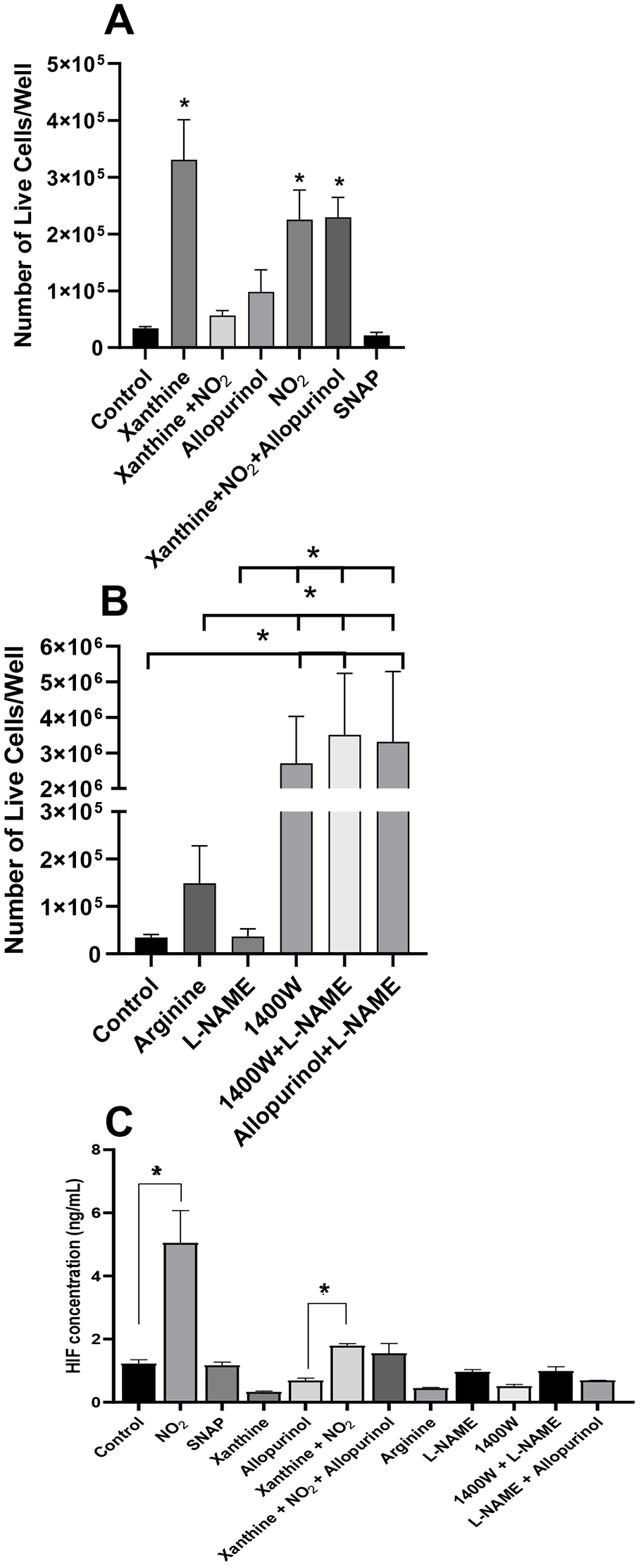
Anaerobic HeLa cell viability modulation by mediators of the nitric oxide generating pathways. (**A,B**) Number of viable cells/well present after 10 days of anaerobic incubation with various mediators of the pathways responsible for nitric oxide generation. Treatments (Mean ± SEM; n=6, repeated once to twice); * = *p* < 0.05 considered significant; (**C**) Hypoxia inducible factor (HIF) expression by HeLa cells grown anaerobically in the presence of various mediators which promote, or inhibit, the generation of nitric oxide. Each HIF measurement is a pooled sample from viability measurements. HIF protein level (Mean ± SEM; n=6, repeated once to twice); * = *p* < 0.05 considered significant.

Like NO generation, multiple parallel pathways are involved in H_2_S synthesis from the precursor cysteine (Fu et al., 2012). These include the cystathionine-β-synthase (CBS) branch, located in the central nervous system; the cystathionine γ-lyase (CSE) branch, located in the vasculature, liver, and kidney; and the 3-mercaptopyruvate sulfur transferase (3-MST) branch, which is found in kidney, liver, lung, heart, muscle, spleen, and brain (Nagahara et al., 1998). H_2_S oxidation, which occurs widely throughout eukaryotes, is coupled to ATP synthesis by a mitochondrial electron transport chain through a sulfide quinone oxidoreductase (Giuffrè and Vicente, 2018; Yong and Searcy, 2001). H_2_S also regulates expression of cytochrome c oxidase (Guo et al., 2012; Murphy et al., 2019; Vicente et al., 2016). Interestingly, the cysteine/CSE/H_2_S pathway is involved in melanoma progression, and has been shown to promote cell cycle progression in squamous cell (oral) carcinoma cells *in vitro* (Bhattacharyya et al., 2013; Chakraborty et al., 2018; Ma et al., 2015; Panza et al., 2014).

Overall, any treatment that increased cellular H_2_S levels, particularly the slow H_2_S donor GYY4137, or the inhibition of CSE, which blocked glutathione, a consumer of H_2_S synthesis, caused significantly (*p* < 0.05) enhanced viability, as compared to medium alone (cysteine, 9.8-; JK-2, 8.8-; and GYY4137, 40.4-fold increases; Fig 2A). This finding is in contrast to reported negative effects on viability when CSE expression (siRNA) is silenced in the presence of oxygen, or after exposure to an oxidizing agent (hydrogen peroxide) (Lee et al., 2014). Since under anaerobic culture there is minimal need for the antioxidative properties of H_2_S to combat cellular reactive oxygen species generated oxidative stress, the predictably higher H_2_S levels could be used to drive mitochondrial ATP production (Fu et al., 2012; Plotkin et al., 2018a; Szabo et al., 2013). Interestingly, except for nitrite which has been reported to affect HIF expression, HIF expression lacked correlation with cell viability under anaerobic growth conditions. In addition, H_2_S had no significant effect on HIF expression, contrary to its effect in *Caenorhabditis elegans* (Fig 1C; 2C) (Budde and Roth, 2010). This finding casts doubt on HIF’s relevance for cells growing long-term anaerobically, in contrast to its importance in hypoxically-grown cells (Singh et al., 2012).

**Fig. 2.**
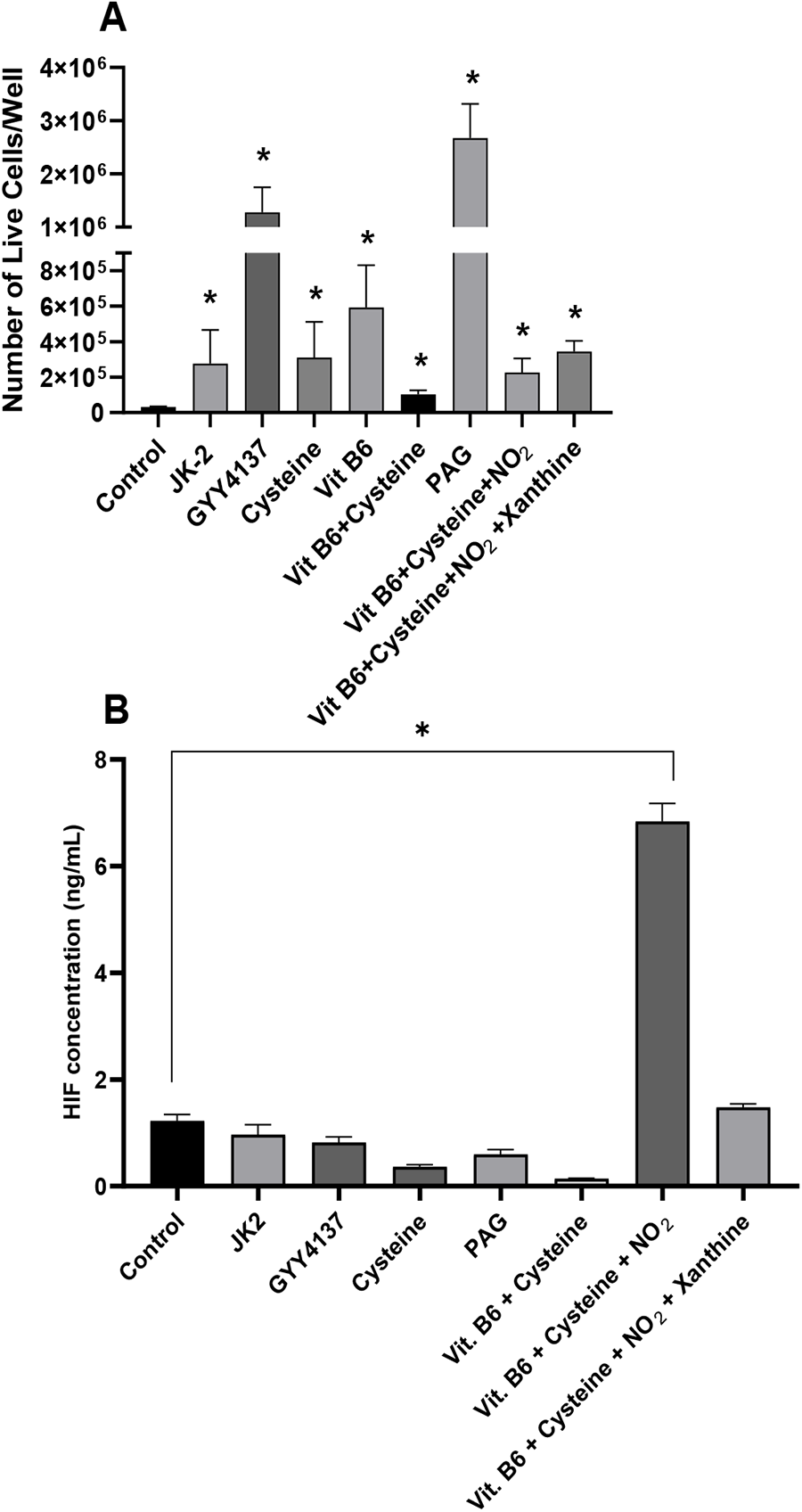
Anaerobic HeLa cell viability modulation by mediators of the hydrogen sulfide pathways. (**A**) Number of viable cells/well present after 10 days of incubation with various mediators of the hydrogen sulfide pathways under anaerobic conditions; (**B**) Expression of hypoxia inducible factor (HIF) response to the various mediators after growth under anaerobic conditions. Each HIF measurement is a pooled sample from viability measurements. Mediators and HIF (Mean ± SEM; n=6, repeated once to twice); * = *p* < 0.05 considered significantly different from control.

Depending on the organ system, and type of NO synthase, H_2_S exhibits contradictory effects. These effects include inhibition of NO produced by the neuronal form of nitric oxide synthase (nNOS), or increased NO production by iNOS and eNOS (Heine et al., 2015; Hou et al., 2017). In addition, H_2_S appears to function as a feedback inhibitor of iNOS and nNOS, presumably due to its function as a free radical scavenger (Kolluru et al., 2013; Kolluru et al., 2015). To determine if constituents of H_2_S producing pathways support HeLa cell anaerobic viability, cysteine, H_2_S donors JK-2 (short term) and GYY4137 (slow release), pathway cofactor pyridoxal phosphate (vitamin B6), or propargylglycine (PAG), the irreversible inhibitor of cystathionine γ-lyase and subsequent glutathione synthesis, were tested (Fig. 2A).

All H_2_S-related supplements tested increased viability (Fig 2A). The most potent enhancers of cell viability were the H_2_S donors GYY4137 and PAG (Diwakar and Ravindranath, 2007; Fu et al., 2012; Lee et al., 2011). The effects for cysteine and vitamin B6 ranged from 10- and 19-fold increase, respectively. As was measured for nitric oxide pathways, nitrite was the sole supplement responsible for increasing HIF expression compared to control (Fig 2B). Interestingly, the level of HIF in cells grown with cysteine alone and with its cofactor vitamin B6 was 30% and 11% that of cells grown in control medium, respectively, indicating that factors that lead to H_2_S production further destabilize HIF protein in cells incubated for extended periods (10 days) in the absence of oxygen. This is the first report of the effects various gaseous chemical signaling molecules, and their inhibitors, have on the viability of anaerobically grown cells, and the role H_2_S plays in sustaining viability in the absence of oxygen. Further studies on anaerobic mitochondrial function are ongoing.

## Author Contributions

Conceptualization, B.P. and I.S..; methodology, B.P. and I.S.; validation, B.P, I.S., and A.K.; formal analysis, B.P, I.S., and A.K.; investigation, B.P, I.S., and A.K.; resources, B.P.; data curation, B.P. and A.K.; writing—original draft preparation, B.P.; writing—review and editing, I.S. and A.K.; visualization, B.P. and A.K; supervision, B.P. and I.S.; project administration, B.P.; funding acquisition, B.P. All authors have read and agreed to the published version of the manuscript.

## Acknowledgments

B.P. thanks Prasanth Puthanveetil, Ph.D. for his helpful discussions.

## References

Bhattacharyya, S., Saha, S., Giri, K., Lanza, I. R., Nair, K. S., Jennings, N. B., Rodriguez-Aguayo, C., Lopez-Berestein, G., Basal, E., Weaver, A. L. et al. (2013). Cystathionine beta-synthase (CBS) contributes to advanced ovarian cancer progression and drug resistance. PLoS One 8, e79167.

Budde, M. W. and Roth, M. B. (2010). Hydrogen Sulfide Increases Hypoxia-inducible Factor-1 Activity Independently of von Hippel–Lindau Tumor Suppressor-1 in C. elegans. Molecular Biology of the Cell 21, 212–217.

Cantu-Medellin, N. and Kelley, E. E. (2013). Xanthine Oxidoreductase-Catalyzed Reduction of Nitrite to Nitric Oxide: Insights Regarding Where, When and How. Nitric oxide: biology and chemistry / official journal of the Nitric Oxide Society 34, 19–26.

Chakraborty, P. K., Murphy, B., Mustafi, S. B., Dey, A., Xiong, X., Rao, G., Naz, S., Zhang, M., Yang, D. and Dhanasekaran, D. N. (2018). Cystathionine β-synthase regulates mitochondrial morphogenesis in ovarian cancer. The FASEB Journal, fj-201701095R.

Chen, S., Yue, T., Huang, Z., Zhu, J., Bu, D., Wang, X., Pan, Y., Liu, Y. and Wang, P. (2019). Inhibition of hydrogen sulfide synthesis reverses acquired resistance to 5-FU through miR-215-5p-EREG/TYMS axis in colon cancer cells. Cancer Letters 466, 49–60.

Cheng, H., Wang, L., Mollica, M., Re, A. T., Wu, S. and Zuo, L. (2014). Nitric oxide in cancer metastasis. Cancer Letters 353, 1–7.

Cojoc, M., Mabert, K., Muders, M. H. and Dubrovska, A. (2015). A role for cancer stem cells in therapy resistance: Cellular and molecular mechanisms. Seminars in Cancer Biology 31, 16–27.

Cummins, J. and Tangney, M. (2013). Bacteria and tumours: causative agents or opportunistic inhabitants? Infectious agents and cancer 8, 11.

Damacena-Angelis, C., Oliveira-Paula, G. H., Pinheiro, L. C., Crevelin, E. J., Portella, R. L., Moraes, L. A. B. and Tanus-Santos, J. E. (2017). Nitrate decreases xanthine oxidoreductase-mediated nitrite reductase activity and attenuates vascular and blood pressure responses to nitrite. Redox Biology 12, 291–299.

De Vries, S. and Schröder, I. (2002). Comparison between the nitric oxide reductase family and its aerobic relatives, the cytochrome oxidases. Biochemical Society Transactions 30, 662–667.

Diwakar, L. and Ravindranath, V. (2007). Inhibition of cystathionine-γ-lyase leads to loss of glutathione and aggravation of mitochondrial dysfunction mediated by excitatory amino acid in the CNS. Neurochemistry International 50, 418–426.

Fu, M., Zhang, W., Wu, L., Yang, G., Li, H. and Wang, R. (2012). Hydrogen sulfide metabolism in mitochondria and its regulatory role in energy production. Proceedings of the National Academy of Sciences 109, 2943.

Giuffrè, A. and Vicente, J. B. (2018). Hydrogen Sulfide Biochemistry and Interplay with Other Gaseous Mediators in Mammalian Physiology. Oxidative Medicine and Cellular Longevity 2018.

Gladwin, M. T., Schechter, A. N., Kim-Shapiro, D. B., Patel, R. P., Hogg, N., Shiva, S., Cannon Iii, R. O., Kelm, M., Wink, D. A., Espey, M. G. et al. (2005). The emerging biology of the nitrite anion. Nature Chemical Biology 1, 308–314.

Gronroos, T. J., Lehtio, K., Soderstrom, K. O., Kronqvist, P., Laine, J., Eskola, O., Viljanen, T., Grenman, R., Solin, O. and Minn, H. (2014). Hypoxia, blood flow and metabolism in squamous-cell carcinoma of the head and neck: correlations between multiple immunohistochemical parameters and PET. BMC Cancer 14, 876.

Guo, W., Kan, J.-t., Cheng, Z.-y., Chen, J.-f., Shen, Y.-q., Xu, J., Wu, D. and Zhu, Y.-z. (2012). Hydrogen sulfide as an endogenous modulator in mitochondria and mitochondria dysfunction. Oxidative Medicine and Cellular Longevity 2012.

Heine, C. L., Schmidt, R., Geckl, K., Schrammel, A., Gesslbauer, B., Schmidt, K., Mayer, B. and Gorren, A. C. (2015). Selective Irreversible Inhibition of Neuronal and Inducible Nitric-oxide Synthase in the Combined Presence of Hydrogen Sulfide and Nitric Oxide. J Biol Chem 290, 24932–44.

Hou, X., Yuan, Y., Sheng, Y., Yuan, B., Wang, Y., Zheng, J., Liu, C. F., Zhang, X. and Hu, L. F. (2017). GYY4137, an H_2_S Slow-Releasing Donor, Prevents Nitrative Stress and alpha-Synuclein Nitration in an MPTP Mouse Model of Parkinson’s Disease. Front Pharmacol 8, 741.

Ivanovic, Z. (2009). Hypoxia or in situ normoxia: The stem cell paradigm. Journal Of Cellular Physiology 219, 271–275.

Kolluru, G. K., Shen, X. and Kevil, C. G. (2013). A tale of two gases: NO and H_2_S, foes or friends for life? Redox Biology 1, 313–318.

Kolluru, G. K., Yuan, S., Shen, X. and Kevil, C. G. (2015). Chapter Fifteen - H_2_S Regulation of Nitric Oxide Metabolism. In Methods in Enzymology, vol. 554 eds. E. Cadenas and L. Packer), pp. 271–297: Academic Press.

Lee, Z. W., Low, Y. L., Huang, S., Wang, T. and Deng, L. W. (2014). The cystathionine γ-lyase/hydrogen sulfide system maintains cellular glutathione status. Biochem J 460, 425–35.

Lee, Z. W., Zhou, J., Chen, C.-S., Zhao, Y., Tan, C.-H., Li, L., Moore, P. K. and Deng, L.-W. (2011). The slow-releasing hydrogen sulfide donor, GYY4137, exhibits novel anti-cancer effects in vitro and in vivo. PLoS One 6, e21077.

Ma, Z., Bi, Q. and Wang, Y. (2015). Hydrogen sulfide accelerates cell cycle progression in oral squamous cell carcinoma cell lines. Oral diseases 21, 156–162.

Murphy, B., Bhattacharya, R. and Mukherjee, P. (2019). Hydrogen sulfide signaling in mitochondria and disease. The FASEB Journal 33, 13098–13125.

Nagahara, N., Ito, T., Kitamura, H. and Nishino, T. (1998). Tissue and subcellular distribution of mercaptopyruvate sulfurtransferase in the rat: confocal laser fluorescence and immunoelectron microscopic studies combined with biochemical analysis. Histochem Cell Biol 110, 243–50.

Pal, V. K., Bandyopadhyay, P. and Singh, A. (2018). Hydrogen sulfide in physiology and pathogenesis of bacteria and viruses. IUBMB Life 70, 393–410.

Panza, E., De Cicco, P., Armogida, C., Scognamiglio, G., Gigantino, V., Botti, G., Germano, D., Napolitano, M., Papapetropoulos, A., Bucci, M. et al. (2014). Role of the cystathionine γ lyase/hydrogen sulfide pathway in human melanoma progression. Pigment Cell & Melanoma Research 28, 61–72.

Plotkin, B. J., Davis, J. W., Strizzi, L., Lee, P., Christoffersen-Cebi, J., Kacmar, J., Rivero, O. J., Elsayed, N., Zanghi, N., Ito, B. et al. (2018a). A method for the long-term cultivation of mammalian cells in the absence of oxygen: Characterization of cell replication, hypoxia-inducible factor expression and reactive oxygen species production. Tissue and Cell 50, 59–68.

Plotkin, B. J., Hatakeyama, T. and Ma, Z. (2015). Antimicrobial susceptibility and sub-mic biofilm formation of *Moraxella catarrhalis* clinical isolates under anaerobic conditions. Advances in Microbiology Vol. 05 No.04, 8.

Plotkin, B. J., Sigar, I. M., Swartzendruber, J. A., Kaminski, A. and Davis, J. (2018b). Differential expression of cytokines and receptor expression during anoxic growth. BMC Research Notes 11, 406.

Singh, M., Arya, A., Kumar, R., Bhargava, K. and Sethy, N. K. (2012). Dietary nitrite attenuates oxidative stress and activates antioxidant genes in rat heart during hypobaric hypoxia. Nitric Oxide 26, 61–73.

Siva Raju, K., Sharma, N. D. and Lodha, M. L. (1997). ATP production during dissimilatory nitrate reduction in Azorhizobium caulinodans IRBG 46. FEMS Microbiology Letters 151, 17–21.

Stoimenova, M., Igamberdiev, A. U., Gupta, K. J. and Hill, R. D. (2007). Nitrite-driven anaerobic ATP synthesis in barley and rice root mitochondria. Planta 226, 465–474.

Szabo, C., Ransy, C., Módis, K., Andriamihaja, M., Murghes, B., Coletta, C., Olah, G., Yanagi, K. and Bouillaud, F. (2013). Regulation of mitochondrial bioenergetic function by hydrogen sulfide. Part I. Biochemical and physiological mechanisms. British journal of pharmacology 171, 2099–2122.

Trageser, M. and Unden, G. (1989). Role of cysteine residues and of metal ions in the regulatory functioning of FNR, the transcriptional regulator of anaerobic respiration in Escherichia coli. Molecular Microbiology 3, 593–599.

Vicente, J. B., Malagrinò, F., Arese, M., Forte, E., Sarti, P. and Giuffrè, A. (2016). Bioenergetic relevance of hydrogen sulfide and the interplay between gasotransmitters at human cystathionine β-synthase. Biochimica et Biophysica Acta (BBA)-Bioenergetics 1857, 1127–1138.

Visvader, J. E. and Lindeman, G. J. (2008). Cancer stem cells in solid tumours: accumulating evidence and unresolved questions. Nature Reviews Cancer 8, 755.

Vlaski-Lafarge, M., Labat, V., Brandy, A., Refeyton, A., Duchez, P., Rodriguez, L., Gibson, N., Brunet de la Grange, P. and Ivanovic, Z. (2020). Normal Hematopoetic Stem and Progenitor Cells Can Exhibit Metabolic Flexibility Similar to Cancer Cells. Frontiers in Oncology 10, 713.

Yong, R. and Searcy, D. G. (2001). Sulfide oxidation coupled to ATP synthesis in chicken liver mitochondria. Comparative Biochemistry and Physiology Part B: Biochemistry and Molecular Biology 129, 129–137.

